# Network influence determines the impact of cortical ensembles on stimulus detection

**DOI:** 10.1101/2024.08.18.608496

**Authors:** Hayley A. Bounds, Hillel Adesnik

**Affiliations:** Department of Neuroscience, University of California, Berkeley; The Helen Wills Neuroscience Institute

## Abstract

Observation of neural firing patterns can constrain theories for the types of activity patterns that the brain uses to guide behavior. However, directly perturbing these patterns, ideally with great specificity, is required to causally test any particular theory. We combined two-photon imaging and cellular resolution optogenetic photo-stimulation to causally test how neural activity in the mouse visual cortex is read out to detect visual stimuli. Contrary to expectations, targeted activation of highly sensitive neural ensembles did not preferentially modify behavior compared to random ensembles, contradicting a longstanding hypothesis for how neural activity drives stimulus detection. Instead, the main predictor of a targeted neural ensemble’s impact on perception was its effect on network activity. This argues that downstream regions summate visual cortex activity without preferentially weighting more informative neurons to make sensory detection decisions. Comparing mouse behavioral performance to decoding models of neural activity implies that mice employ this simple, albeit suboptimal strategy to solve the task. This work challenges conventional notions for how sensory representations mediate perception and demonstrates that specific neural perturbations are critical for determining which features of neural activity drive behavior.

## Introduction

Detecting stimuli is an essential function of sensory systems. Yet behaviorally relevant stimuli are frequently weak, and natural environments are often complex and noisy, complicating detection. In the sensory neocortex, decades of observational data have identified neurons that encode specific stimuli with high sensitivity^1–10^. This work has led to the commonly-held theory that the circuits downstream of the sensory cortex preferentially weight these highly informative neurons to drive perceptual decisions^1,2,11–16^. These and other features of the neural code could be read out, or decoded, in multiple ways to determine the presence of stimuli. While observational study can constrain the types of computational strategies downstream circuits can use to drive behavior, causal manipulations are required to conclusively distinguish which strategies the brain uses.

To test longstanding theories about the cortical codes of sensory detection, we sought to determine which features of neural activity in the mouse primary visual cortex (V1) causally drive perceptual decisions in a visual contrast detection task. Many studies have employed relatively non-specific perturbations, such as lesions or wide-field optogenetic suppression, to probe the role of V1 in sensory behaviors. This work has demonstrated that V1 is critical for a wide array of visual behaviors, including contrast detection, orientation discrimination, change detection, and others^17–22^. Conversely, activation of V1 is perceptually detectable: cortical stimulation in humans generates phosphenes, and broad activation of V1 is sufficient to induce responses in animals trained on detection tasks^19,23–25^. More precise photostimulation in the primary sensory cortex can also impact behavior: high resolution 2-photon photo-stimulation^26–29^ is both detectable in artificial detection tasks^30,31^ and can impact performance in visual discrimination^32,33^ and detection^34^. However, none of these studies determined whether stimulating neural ensembles with different visual response properties differentially impacts detection. Thus, the specific coding strategy used for visual detection remains unknown.

In V1, contrast is a fundamental coding variable for stimulus intensity with high contrast driving more overall cortical activity^7,19,21,35,36^. Thus, to detect visual stimuli, downstream circuits could equally weight the activity of all V1 neurons, regardless of their sensitivity to the stimulus^19,21,37,38^. If this were true, the effects of targeted photostimulation should not depend on the visual sensitivity of the stimulated neurons. Alternatively, due to the heterogeneity in how V1 neurons encode stimulus information^19,36,39^, the downstream decision circuit might preferentially weight the activity of V1 neurons based on their responses to a particular stimulus. If this hypothesis were true, targeted photo-stimulation of responsive neurons should be particularly effective at inducing behavioral responses. In support of this theory, many observational and theoretical studies of neural activity in detection and discrimination have suggested that high sensitivity neurons play a privileged role in optimal readout^2,11–14,40–42^. Indeed, the notion that brain circuits weight neurons that carry more information to drive behavior is arguably one of the most intuitive ideas we have about neural codes.

Even while V1 projection neurons show heterogenous responses to visual stimuli, they also exert heterogenous effects on their local network through recurrent circuits^43^. Indeed, their recurrent impacts onto other projection neurons within V1 may be just as critical for determining how they ultimately influence behavior whether the downstream readout uses a weighted or an unweighted decoding scheme. However, it remains unknown how the sign and magnitude of an ensemble’s recurrent impact influences its impact on behavior.

To address these questions, we imaged and perturbed activity at high resolution in Layer 2/3 V1 excitatory neurons during performance of a contrast detection task. While a weighted decoder more accurately predicted stimulus presence than an unweighted (mean) decoder, neither fully captured the mouse’s behavior. Thus, imaging alone was insufficient to conclusively differentiate between these possibilities. Next, we used high resolution 2-photon holographic photo-stimulation to specifically activate different neural ensembles during task performance. Contrary to long standing assumptions, the visual stimulus sensitivity of the stimulated neurons did not predict their impact on behavioral performance, arguing against the concept that these neurons play a privileged role in behavioral readout. Instead, the main predictors of a targeted ensemble’s impact on stimulus detection were a combination of the activity of the targeted neurons themselves, and more importantly, their recurrent impact on the nearby network. Activating V1 ensembles that facilitated V1 network activity substantially increased stimulus detection, while activating ensembles that reduced V1 output reduced stimulus detection. Our data highlight that the recurrent impact of neurons in V1 are a key determinant of their role in perception. Taken together, our results argue that detection relies on a simple strategy that may facilitate generalization at the expense of optimality: readout of the entire population activity with no privileged role for specific stimulus-sensitive neurons. This suggests that suboptimal performance may not result from readout noise in downstream regions but from suboptimal readout strategies that may be more generalizable in variable natural contexts.

## Results

### Highly sensitive neurons can support visual detection

To test the role of neural activity patterns in stimulus detection, we trained mice to detect a small (20°) visual grating of a fixed orientation^24,44^ (Fig. 1A, S1). Briefly, mice reported stimulus detection in uncued trials by licking in a response window. Licking during catch trials with no visual stimulus presence or intertrial intervals was punished with a timeout. Prior work has demonstrated that V1 neural activity is necessary for performance in this and similar tasks^18–20,44^. We chose a relatively small stimulus so that the majority of the retinotopically-aligned, visually-driven activity would be within our imaging field of view, allowing access to a large fraction of putatively relevant neurons in L2/3^45^. We expressed GCaMP6s in excitatory pyramidal cells and imaged multiple planes of neural activity simultaneously in V1 L2/3 of trained mice with 2-photon calcium imaging during task performance.

**Figure 1:**
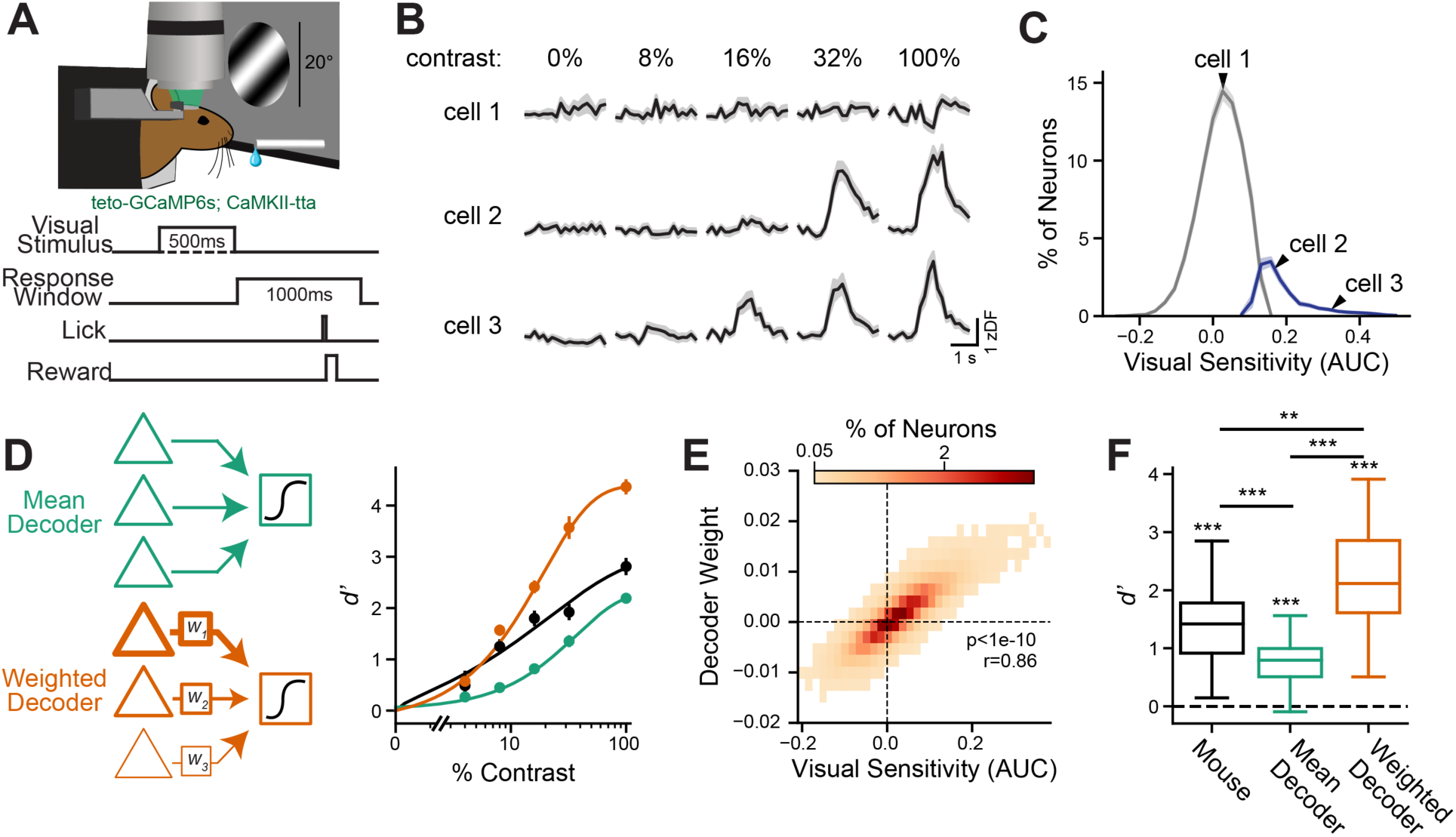
V1 neural activity in a contrast detection task and decoding strategies. **A:** Schematic of the visual detection task where in uncued trials a static, 20° grating was shown for 500 ms, followed by a 1000 ms response window where the mouse had to lick to receive a water reward if a stimulus was present but otherwise withhold licking. Activity was measured with 2-photon imaging of GCaMP6s. **B:** Three example cells with increasing visual sensitivity. **C:** Mean distribution of visual sensitivity (calculated by ROC analysis) of significantly sensitive cells (blue) and all other cells (gray), over 31 sessions in 12 mice. Arrows indicate where each example cell from B falls on the curve based on its sensitivity. AUC: area under the ROC curve. **D:** Comparison of the *d’* (discriminability index) of mouse behavior (black), and two linear classifiers trained to detect stimulus presence. Teal, a decoder on the mean activity of all imaged neurons. Orange, a decoder allowed to vary the weights of individual neurons. **E:** Decoding weight in the weighted decoder vs visual sensitivity for all neurons, displayed as a 2D histogram. Colorbar, % of neurons in each bin. Statistics are for Pearson’s correlation coefficient; n=30,141 neurons from 31 sessions. **F:** Comparison of cross-validated *d’* of different models tested at test contrasts (contrasts between 1-99%). Difference from chance: mouse: p=7.0e-6, mean decoder: p=8.6e-6, weighted decoder: p=7.0e-6. Between models: mouse vs mean decoder p=3.8e-4, mouse vs weighted decoder p=8.4e-3; mean decoder vs weighted decoder p=7.0e-6 multiple-comparisons corrected Wilcoxon signed-rank test. Data plotted as mean ± s.e.m. unless otherwise noted.

We first assessed the responses of V1 pyramidal neurons to the task stimulus. To reduce confounds of movement-related activity, we only considered activity during the ∼650 ms after visual stimulus onset, which was prior to lick responses in most cases (Fig. S1B). We used receiver operating characteristic (ROC) analysis to measure how well the activity of an individual neuron can detect stimulus presence on a trial-by-trial basis^40,46,47^. We defined ‘visual sensitivity’ as the area under the ROC curve (AUC), and it could vary from −0.5 to 0.5, where 0 reflects chance performance, 0.5 is perfect positive prediction (i.e., increased activity predicts stimulus presence) and −0.5 is perfect negative prediction (i.e., decreased activity predicts stimulus presence). This metric measures detectability across all contrasts, so it depends on the strength and variability of a neuron’s visual response as well as on its contrast threshold. On average, 15 ± 1% of the imaged L2/3 pyramidal neurons had significant positive visual sensitivity (calculated by bootstrapped confidence intervals, n=31 sessions in 12 mice). Within these neurons, there was substantial variability in sensitivity (Fig. 1B-C).

The relatively small fraction of highly sensitive neurons suggests that the animal might benefit from a weighted readout strategy to preferentially integrate activity from these highly selective neurons. However, we can also consider an alternative strategy where the downstream circuit weights all inputs equally and simply summates all afferent activity (Fig. 1D). While this latter strategy might seem suboptimal, it might benefit from being more generalizable across stimuli. We explored models reflecting each of these scenarios. These models are not meant to represent the specific biological mechanism of decision making or the most optimal performance strategy, but rather to gain intuitions about possible neural coding schemes. Using ROC analysis, we found that a logistic regression classifier trained on the mean of all observed neural activity, without preferentially weighting any neurons, could reliably detect the presence of the visual stimulus at test contrasts, but performed significantly worse than the animals (Fig. 1E-F, mouse vs mean decoder: p=3.8e-4, mean decoder vs chance: p=8.6e-6). This difference could indicate that mice use a more sophisticated decoding strategy. However, since our estimate of mean activity is limited by only imaging a subpopulation of the total visually driven cortical population, as well as by imperfections in activity estimation via calcium imaging^48^, it is still possible that a mean readout of all V1 activity might be a suitable strategy for the mouse.

We next examined the performance of a logistic regression model allowed to vary weights across neurons (Fig. 1D). When trained to detect stimulus presence, this decoder weighted neurons with high visual sensitivity more strongly than those with low sensitivity (Fig. 1E). The weighted decoder performed substantially better at detecting the stimulus than the mean activity decoder (Fig. 1F, weighted decoder vs mean decoder: p=7e-6). Surprisingly, the weighted decoder performed even better than the mouse did at detecting the stimulus, suggesting that the information present in V1 could theoretically support better performance, consistent with other work (Fig. 1F, mouse vs weighted decoder: p=0.0084)^7,49^. While these analyses suggest that a weighted strategy is optimal and may account better for the performance of the mouse, these observational approaches are insufficient to conclusively determine what decoding strategy the mouse brain uses to detect visual stimuli. Therefore, we sought to test what strategy the mouse brain uses to detect visual stimuli by directly altering neural responses in V1.

### Targeted 2-photon photostimulation drives increased V1 activity

To differentiate between possible codes for visual detection, we sought to stimulate ensembles with varying properties. By varying the visual sensitivity and the size of the ensemble, we could assess whether neurons with high sensitivity are weighted more strongly or if total activity drives stimulus detection. To achieve this, we used mice expressing ChroME2s, a powerful excitatory opsin, and GCaMP6s in excitatory pyramidal neurons and performed targeted 2-photon holographic optogenetics in behaving animals in a retinotopically-aligned region of V1^50^ (Fig. 2A). We first assessed the visually driven responses of each neuron in the imaging volume as well as its response to targeted optogenetic stimulation. We then targeted ensembles of neurons that were either visually-sensitive or random. Careful quantification of the effective resolution of our photo-stimulation system demonstrated that it could target light to specific neurons (Fig. 2B, Fig. S3). Based on this, we categorized ‘directly activated’ neurons as those that received direct optogenetic excitation (i.e., were within 21 microns of an illumination spot and that were significantly activated by photostimulation). In contrast, we categorized ‘indirect neurons’ as those that were far enough away (> 21 microns) from any holographic target such that their modulated activity could largely be attributed to synaptic effects through the network, rather than to direct optogenetic excitation. Photo-stimulation groups varied in size (10-48 spots, 7-40 neurons directly activated).

**Figure 2:**
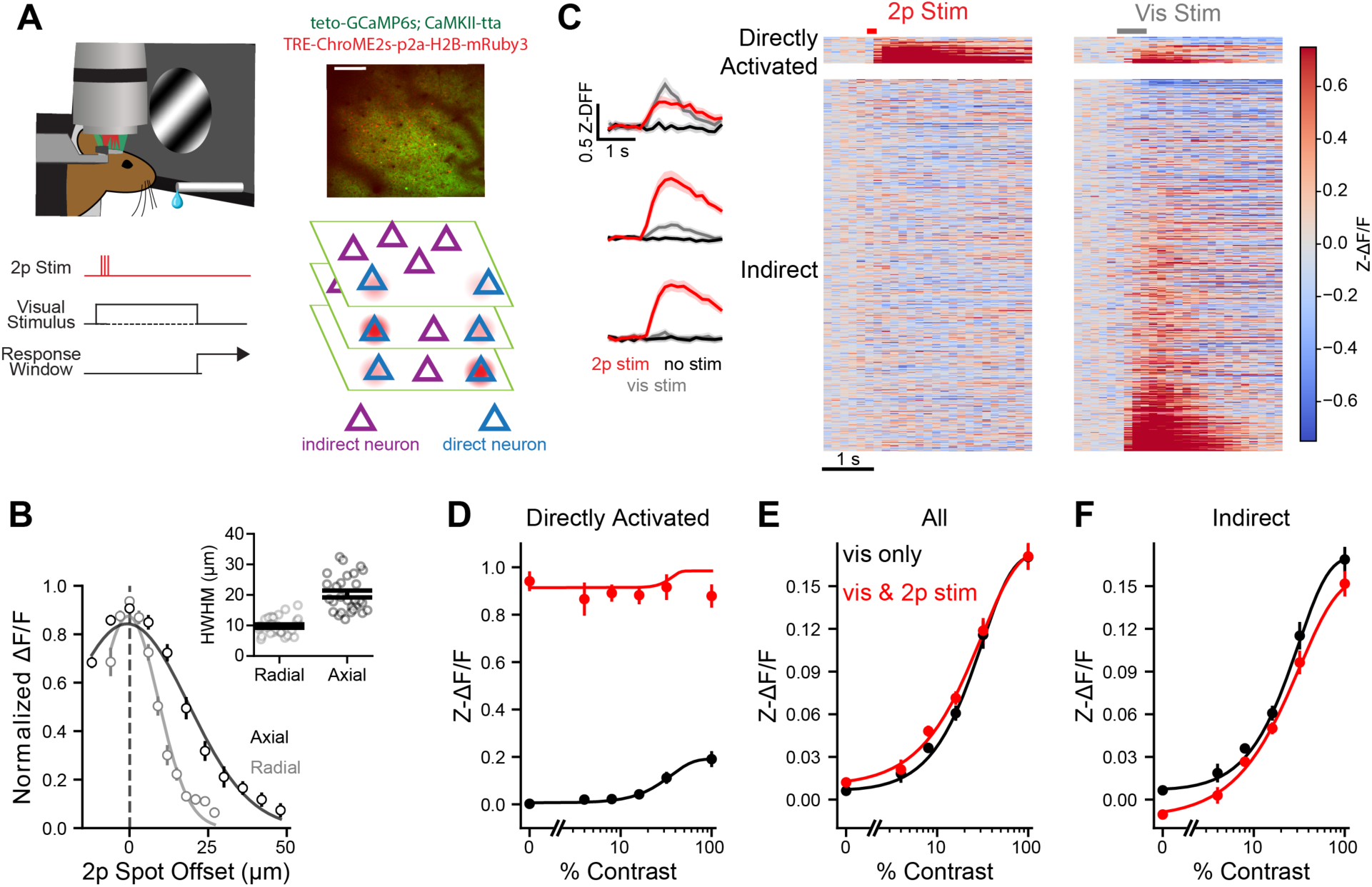
2-photon stimulation of functionally defined cortical ensembles. **A:** Left, experiment schematic. 2-photon single-cell targeted stimulation of neurons during task performance. Stimulation consisted of 3, 5-ms long pulses delivered coincident to the arrival of the first visually-evoked spikes in visual cortex. Right top, example imaging field of view showing GCaMP6s (green) and ChroME2s expression (red). Right bottom, schematic showing distinction between direct neurons (blue) and indirect neurons (purple). Directly activated neurons are activated by 2-photon stimulation spots (red). **B:** Physiological point-spread functions for two-photon activation found by moving the two-photon stimulation spot off-center radially (light gray) and axially (dark gray). Left, normalized response with gaussian fit. Inset: Half-width at half-max (HWHM) of gaussian fits. Points are individual cells. HWHM: radial: 9.7 ± 0.5 μm, axial: 20 ± 1 μm. n=28 neurons in 3 mice. **C:** Photoactivation and visual activation for an example ensemble. Left, mean calcium traces example directly activated neurons for three trial types: no stimulus (black), 100% contrast visual stimulus (gray) and 2-photon stimulation (red). Right, heat maps of directly activated neuron (top) and indirect neuron (bottom) responses during 2-photon stimulation (left) and visual stimulus presentation (right). Red and gray bars, onset and duration of 2-photon and visual stimuli, respectively. Neurons sorted within each map by response strength in held out half of trials, other half plotted here. **D-F:** Responses of directly activated (D), all (E) and indirect (F) neurons across contrasts. Black, visual stimulus only. Red, visual stimulus plus 2-photon stimulation. Effect of photostimulation, two-way repeated measures ANVOA: D: p<1e-5, E: p=0.0007, F: p<1e-5. Data plotted as mean ± s.e.m. unless otherwise noted.

Photostimulation strongly drove activity of directly activated neurons (Fig. 2D, p<1e-5), on average substantially exceeding their visually driven response. Photostimulation increased the mean population response of all neurons in the imaging volume (Fig. 2E, p=0.0088). This increase, however, was driven by the small number of directly activated neurons since the indirect population, on average, showed mean suppression (Fig. 2F, p=0.0003). These data demonstrate that we obtained potent optogenetic control of visual cortical neurons and could significantly alter activity patterns in V1 during the behavioral task.

### An ensemble’s visual sensitivity does not predict its behavioral impact

To discriminate between the two classes of decoding schemes discussed above, we reasoned that we could compare the perceptual consequences of photostimulating visually-sensitive ensembles of neurons that encode the task stimulus as compared to photo-stimulating random ensembles. If the downstream readout preferentially weights neurons that encode more stimulus information (i.e., respond strongly to the stimulus) then photo-stimulating visually sensitive ensembles should more strongly bias behavior than random stimulation. Conversely, if the downstream readout weights all V1 neurons equally, then activating visually sensitive or randomly selected neurons should have similar effects on the animal’s detection performance. We selected ensembles online for these properties, and classified ensembles as visually sensitive or random post-hoc based on their visual sensitivity (Methods). Visually sensitive ensembles responded more to the visual stimulus but were on average matched in size to random ensembles (Fig. 3A-B, p=0.74, n=14 visually sensitive ensembles and 27 random ensembles). These ensemble types did not have different effects on overall network activity (Fig. S4A-B, indirect: p=0.62, all: p=0.74). However, targeted photostimulation of sensitive neurons augmented the total activity of all visually responsive neurons, while targeted photostimulation of random ensembles had no mean effect (Fig. 3C, sensitive ensembles: p=0.0043, random ensembles: p=0.60). These results demonstrate that our photostimulation system is sufficient to execute the intended experiments.

**Figure 3:**
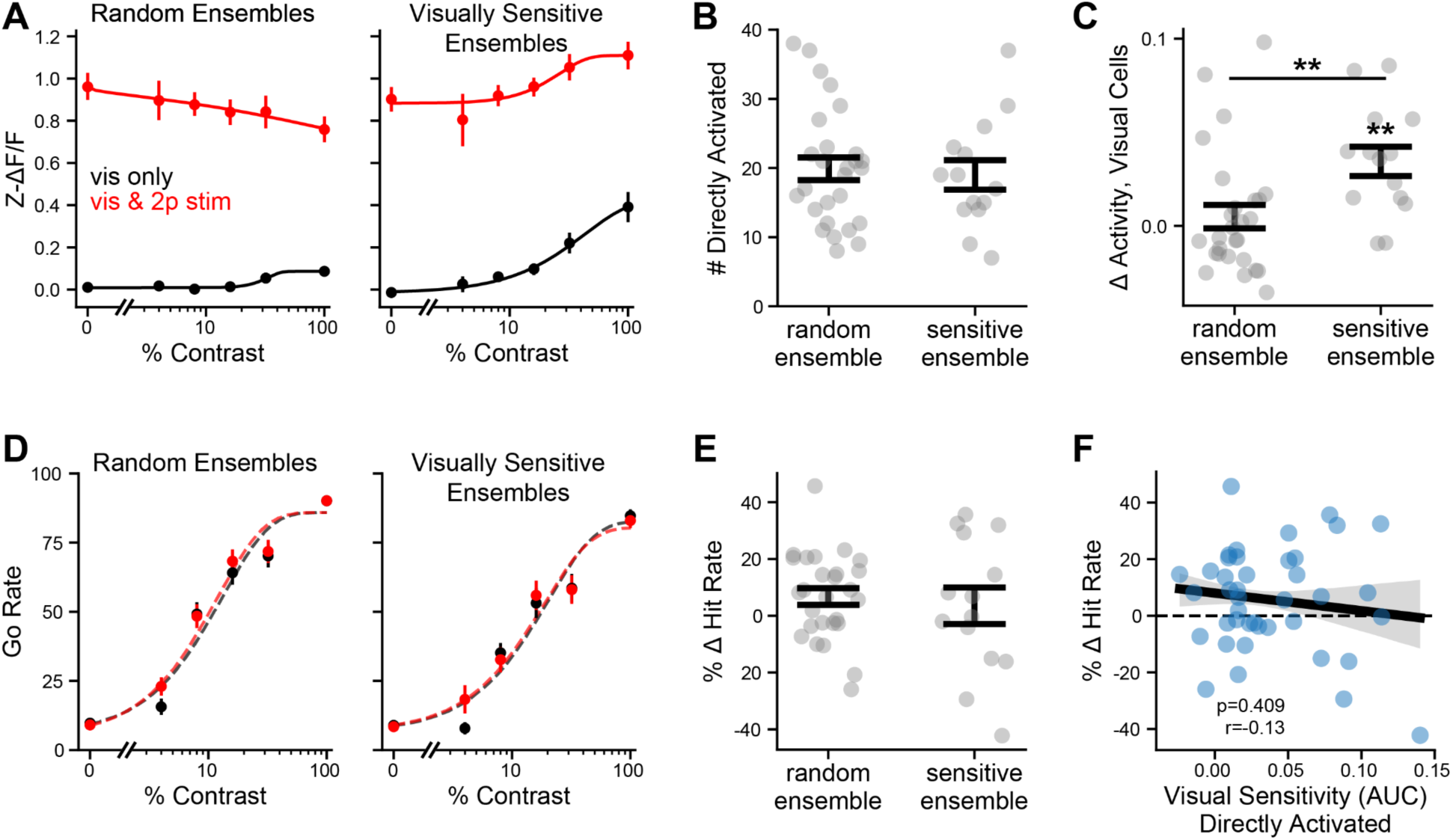
The visual sensitivity of activated neurons does not predict behavioral change. **A:** Visual (black) and visual plus photostimulation (red) responses of random (left) and visually sensitive (right) ensembles, averaged over ensemble. n=27 random and 14 sensitive ensembles. **B:** Number of directly photostimulated neurons in random and sensitive groups. **C:** Change in activity of all (direct and indirect) visually sensitive neurons in no visual stimulus trials, for different ensemble types. Random vs sensitive: p=0.0035 Wilcoxon rank-sums test; within-sensitive stimulation vs no stimulation: p= 0.0019, within-random: p=0.83 Wilcoxon signed-rank tests. **D:** Average psychometric curves for random ensembles (left) and sensitive ensembles (right). Black, visual stimulus only trials. Red, visual stimulus plus 2-photon stimulation. **E:** Change in behavior for different ensemble types. p=0.72 Wilcoxon rank-sum test. **F:** Relationship between mean visual sensitivity (AUC) of directly activated ensemble to behavioral change. Gray area, 68% confidence interval of regression. Statistics for Pearson correlation coefficient. Data are presented as mean ± s.e.m. unless otherwise noted.

Given this, we would expect that if these visually sensitive neurons were weighted more strongly, stimulation of the sensitive ensembles should bias behavior more than random ensembles. Strikingly, there was no difference in the mean effect on behavior between photo-stimulating visually-sensitive vs. random ensembles of neurons (Fig. 3D-E, p=0.98). Further analysis revealed that neither the average visual sensitivity nor the percentage of significantly responsive cells in the directly stimulated population predicted the behavioral change (Fig. 3F, Fig. S4C, p=0.67, n=41 ensembles).

A recent study in barrel cortex suggested that an ensemble’s information about the animal’s choice determines the impact of an ensemble on behavior in a two choice task^46^, and many others have posited that a correlation with choice of the animal suggests that a neuron is involved in downstream readout^1,6,13,37,51–54^ (though see also^37,51^). To examine this possibility, we quantified ‘choice selectivity’, also called ‘choice probability’ – an ROC analysis of the correlation of a neuron’s trial by trial response with the animal’s behavioral choice within matched visual conditions^53,55^. 9 ± 1% of neurons had significant choice selectivity (determined by bootstrapped confidence intervals). Despite seeing above-chance levels of choice information in the population, we found no link between choice coding in directly activated neurons and the behavioral outcome of stimulation (Fig. S4D, p=0.90). These data argue that visually sensitive or choice-coding excitatory neurons in L2/3 of mouse V1 have no privileged role in driving behavioral responses in the contrast detection task, suggesting that downstream regions do not preferentially weight more informative neurons in this circuit.

While we showed above that the visual sensitivity of the directly activated neurons doesn’t predict behavioral performance changes, it is possible that the visual sensitivity of the recurrently activated network is predictive, which would still argue for a weighted readout. Thus, we tested whether the effect of an ensemble might rely on the properties of the putatively synaptically-recruited follower cells, i.e. the significantly activated indirect cells. Analyzing ensembles that had at least four follower neurons (n=34), we found that the visual sensitivity of followers or of all recruited neurons (both photostimulated and indirect followers) did not predict behavior (Fig. S5A). Furthermore, like the directly activated population (Fig. S4C-D), neither the percent of visually responsive neurons in the recruited population nor the choice selectivity of these populations predicted behavior (Fig. S5B-C).

### A cortical ensemble’s network influence predicts its behavioral impact

To test the alternative strategy in which the downstream decoder simply summates V1 activity without weighting specific neurons, we examined the behavioral impact across all photostimulated ensembles. First, we examined the mean effect on performance across all photostimulated ensembles. We found a small but insignificant positive change in behavior at test contrasts (Fig. 4A-B, p=0.054). However, there was considerable heterogeneity in the behavioral impact across targeted ensembles. Indeed, within-session analysis showed that 20% (9/41) of ensembles had a significant effect, a rate higher than expected by chance, indicating that 2-photon optogenetic stimulation of specific ensembles of L2/3 pyramidal neurons can alter stimulus detection (Fig. S6A, significance based on bootstrapped confidence intervals). Photostimulation did not change the false alarm rate, implying that our ensembles might not be large enough to generate a response *de novo* (Fig. S6B-C).

**Figure 4:**
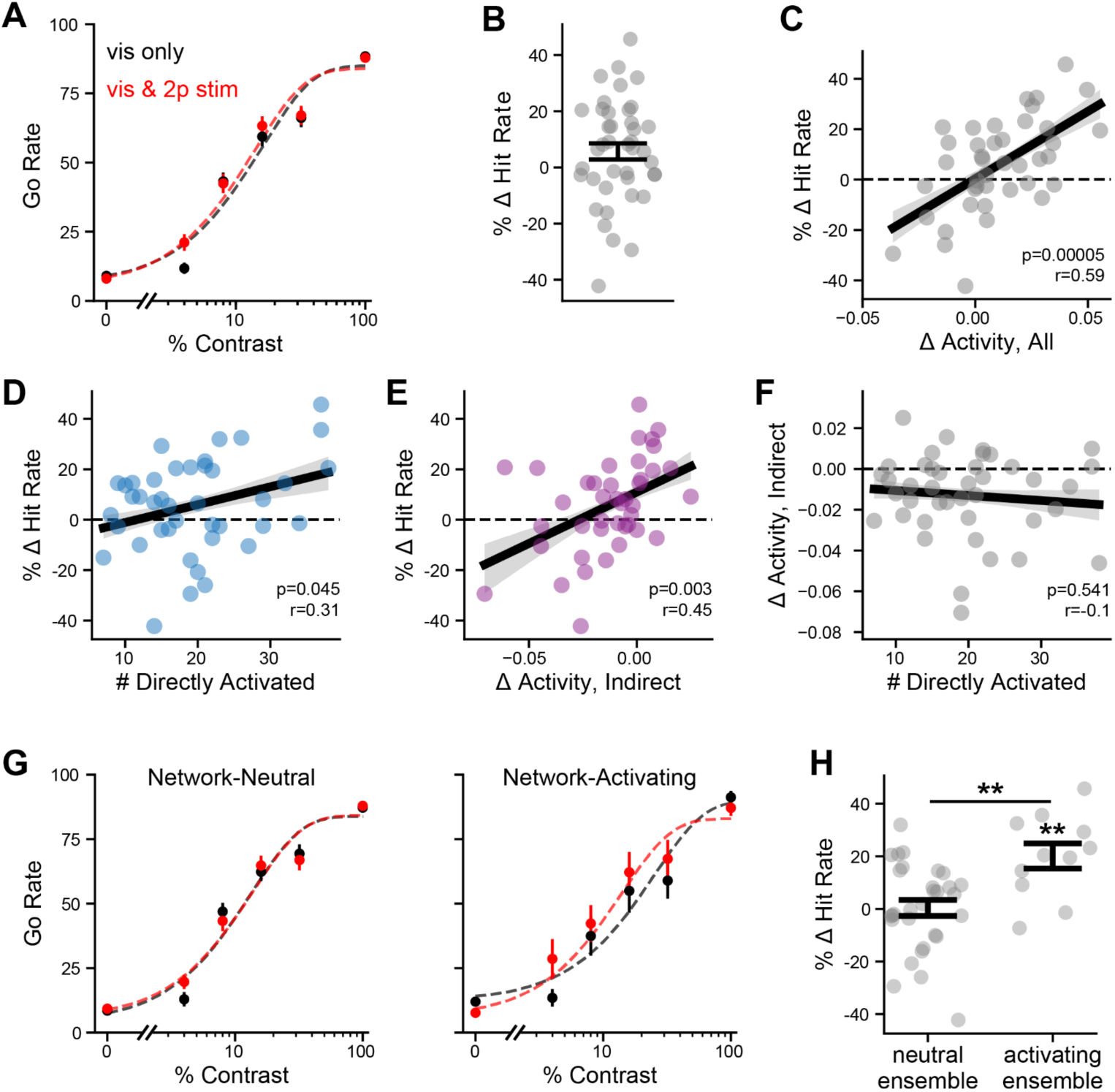
Network activation predicts the behavioral impact of 2-photon holographic stimulation. **A:** Average psychometric curve fit with modified weibull function (dashed lines) across all stimulated ensembles (n=41). Black, visual stimulus only. Red, visual stimulus plus 2-photon stimulation. **B:** % change in hit rate at test contrasts (between 1-99%) between 2-photon stimulation and no stimulation. Gray dots, individual ensembles. p=0.054, Wilcoxon signed-rank. **C:** Relationship between change in neural activity and change in behavior induced by 2-photon stimulation. Statistics are for Pearson’s correlation coefficient. Black line, linear regression with 68% confidence interval shaded. Points, individual ensembles. **D:** As in C, but for relationship between # of directly activated neurons and change in behavior. **E:** As in C, but for the relationship between change in behavior and change in indirect population activity. **F:** As in C, but for relationship between # of directly activated neurons and change in indirect population activity. **G-H:** Difference in behavioral change between neutral ensembles and network-activating ensembles. **G**: Average psychometric curves for each ensemble type. Black, visual stimulus only trials. Red, visual stimulus plus 2-photon stimulation. **H:** % change in hit rate for ensemble types. Neutral vs activating: p=0.0026, Wilcoxon rank-sums. Within neutral, stimulation vs no stimulation: p=0.73, within activating stimulation vs no stimulation: p=0.008, Wilcoxon signed-rank. Data plotted as mean ± s.e.m. unless otherwise noted.

We considered that the substantial heterogeneity in the behavioral impact across photostimulated ensembles should reflect an underlying mechanism that transforms neural activity into stimulus detection. More specifically, if the mouse equally weights V1 neurons in detection, the overall change in network activity a given ensemble generates should predict its behavioral impact. To address this idea, we analyzed each ensemble’s impact on V1 network activity. We restricted analysis to trials with no visual stimulus and no lick responses to avoid the confounds of visual and behavioral signals, such as movement preparation, on network activity^56,57^. This is important because if photostimulation changes the response statistics of the mouse, any change in the network activity would be a combination of local photostimulation-induced changes and changes in the preparatory activity signals, which would be impossible to separate. Hence, we assessed the impact of targeted photostimulation in absence of any other task related signals. In these conditions, the change in total visual cortex activity induced by activation of an ensemble strongly predicted the sign and magnitude of the behavioral change (Fig. 4C, p=5e-5); ensembles that increased total network activity increased performance and vice versa.

This relationship between an ensemble’s network influence and its effect on behavior might be explained by the direct effects on the photostimulated neurons, the recurrent synaptic effects on the indirect population, or both. Although both the number of directly photostimulated neurons and the average activity change in indirect cells predicted the behavioral impact, the recurrent effect on indirect network activity appeared to explain more of the behavioral impact across targeted ensembles (Fig. 4D-E, directly activated: p=0.045, indirect activity: p=0.003). To test the robustness of these results, we also quantified neural activation differently - by the number of significantly recruited cells (‘follower’ cells and total cells) and by the overall change in directly activated neurons. This analysis yielded similar results, demonstrating that our conclusions are robust to different ways of analyzing the neural data (Fig. S7). Together they imply that a neural ensemble’s recurrent impact on total network activity is a key driver of its behavioral impact. Notably, this latter effect was not explained by a correlation between the directly activated activity and the indirect impacts of the ensemble, so these two components could be separated (Fig. 4F, p=0.85). These results thus demonstrate that the recurrent impact of an ensemble is a mechanism by which small perturbations to V1 activity can propagate into changes in behavior. To compare psychometric functions, we divided ensembles into network-neutral (n=30) and network-activating (n=11). Network-activating ensembles had a significant impact on behavior, and a significantly more positive effect on detection rate than network-neutral ensembles (Fig. 4G-H, activating vs neutral: p=0.002, activating vs chance: p=0.008). The importance of both the direct population activation and indirect population activation support that the total activity, regardless of identity, could be used to determine stimulus presence.

### Behavioral changes are consistent with an unweighted readout strategy

To further explore how these findings relate to the possible read out strategy of a downstream region, we returned to the two decoding models we considered earlier. We asked how photo-stimulation altered the ability of decoding models to detect the visual stimulus from the neural data. To this end, we trained decoding models only on non-photostimulation trials and then probed them with neural activity from photostimulation trials. We then compared the change in performance of the decoder as a function of the visual sensitivity of the targeted ensemble (measured in only no visual stimulus, no response trials, Fig. 5A) or the change in network activity the ensemble induced (Fig. 5B). Consistent with the behavior of the mouse, the total network activation but not the visual sensitivity of the stimulated neurons predicted the change in the mean activity decoder’s performance (Fig. 5A-B, visual sensitivity: p=0.289, network activity: p=0.022). Conversely, we found opposite effects of photostimulation for the weighted decoder: unlike the mouse’s behavior, the visual sensitivity of an ensemble strongly predicted the change in the weighted decoder’s performance, while total network activity had no relationship to its performance (Fig. 5C-D, visual sensitivity: p=0.0003, network activity: p=0.174). In addition, across mice, the change in performance of the mean decoder was significantly correlated with change in mouse performance, while the weighted decoder was not (Fig. 5E-F, mean decoder: p=0.02, weighted decoder: p=0.12). These results were similar for other classifier types (Fig. S8). Thus, we found that the impact of photostimulation on the mean activity decoder was significantly more aligned with the impact of photostimulation on mouse behavior than that of the weighted decoder. The fact that holographic photostimulation could augment the performance of the weighted linear classifier but not impact behavioral performance (in the same mice on the same trials) argues against a weighted readout strategy even though it is more optimal for solving the task. Taken together, these experiments support a scheme where the mouse reads out V1 activity in the task with little or no preferential weighting of more visually sensitive neurons.

**Figure 5:**
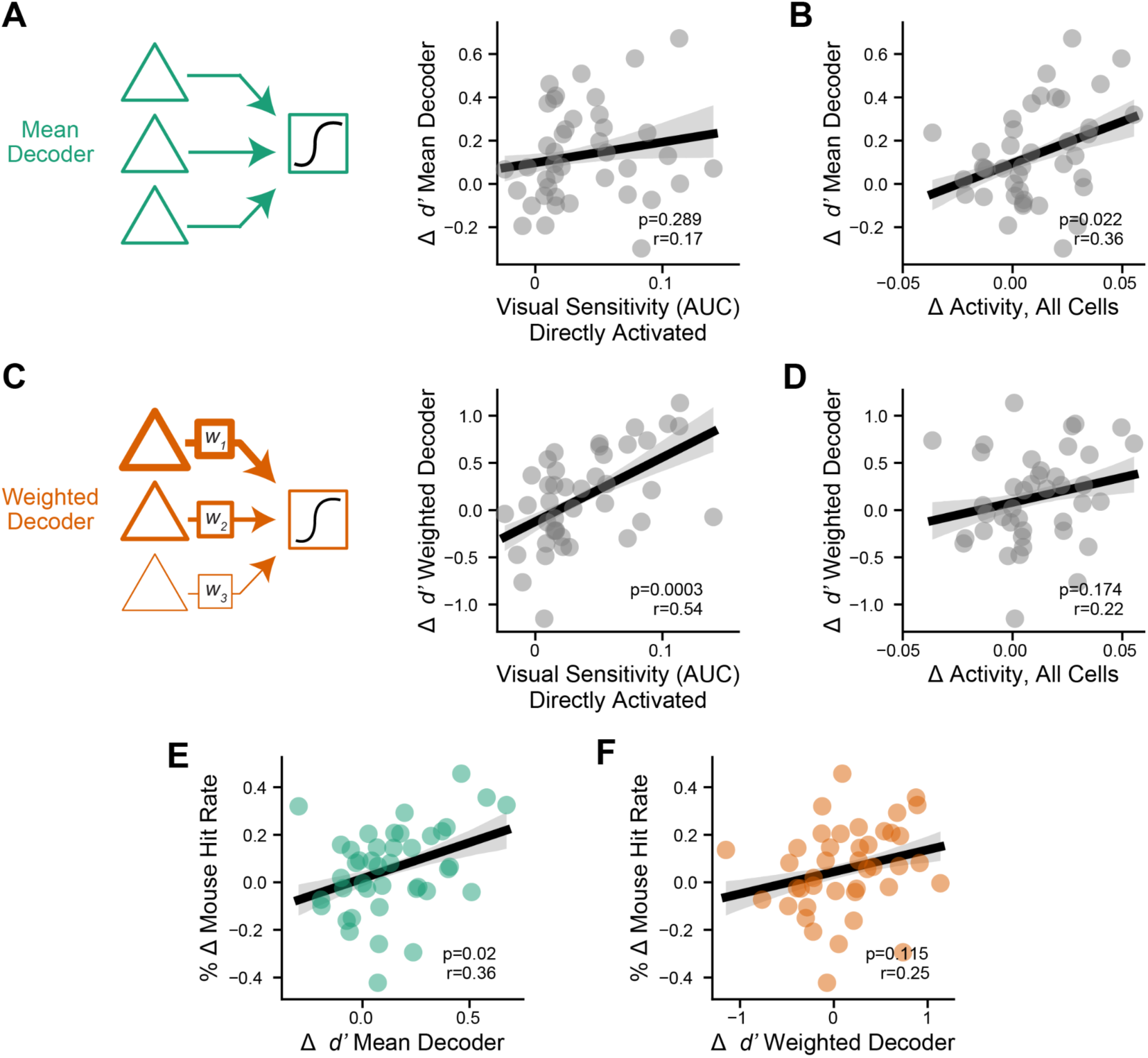
The impact of 2-photon photostimulation on a mean activity decoder is aligned with its impact on behavior. **A:** Change in mean activity decoder *d’* (discriminability index) between 2-photon stimulation and no stimulation trials at test contrasts vs average visual sensitivity (AUC) of directly activated neurons. Gray area, 68% confidence interval of regression. Statistics are for Pearson’s correlation coefficient. **B:** Same as A, but for photostimulation-induced change in activity of all neurons. **C-D:** Same as A-B, but for weighted logistic regression decoder. **E:** Change in mean decoder performance vs mouse performance. **F:** same as E, but for weighted decoder.

## Discussion

Our data imply that mice detect visual stimuli by reading out the mean activity of the visual cortex without preferentially weighting any specific neurons. Although a decoding strategy where mice would preferentially weight more visually sensitive neurons would support higher performance in the task, our targeted photo-stimulation experiments do not support such a scheme. This raises the questions why the mice might use a seemingly suboptimal decoding strategy. One possibility is that an unweighted readout strategy may promote generalization at the cost of optimal performance. Contrast detection is only one of many change detection scenarios that cause a net increase in visual cortex spiking - contrast decrements and orientation changes can also drive overall increased cortical activity, in part due to cortical adaptation to a stimulus (prior to its change)^19,58,59^. This may provide ethological advantages in natural contexts where the specific features of a stimulus are either unknown or irrelevant to the ultimate behavior.

Our results further demonstrate that the recurrent network impact of an ensemble in mouse V1 L2/3, not its visual selectivity, is a key predictor of its impact on behavioral detection of simple visual stimuli. This was not observed in previous studies of detection of artificial photostimulation itself, which depended primarily on the number of directly stimulated neurons but not recurrent impact^30,31^. This difference may be due to several factors including differences in readout strategy, perhaps stemming from the fact that real stimuli, when near detection threshold, tend to activate many neurons weakly, compared to 2-photon stimulation detection, which in recent studies involved the activation of a few neurons very strongly. It may also be due to the fact that our photostimulation was presented coincident with visual stimuli, and the network conditions may thus be quite different. The properties and mechanisms that determine these recurrent impacts, perhaps in a state-dependent fashion, are an important area for future research.

Understanding how V1 activity propagates downstream to areas that summate its output is a logical next step to understand the brain circuits for stimulus detection. Potential areas include the superior colliculus (SC) and various higher visual areas (HVAs), which are necessary for detection and receive strong input from V1^20,60–63^. 2-photon photostimulation combined with 2-photon mesoscale imaging^64^ of HVAs or electrophysiology of SC would enable examination of how information is transformed across regions and how these transformations relate to the behavioral readout strategy. While our evidence suggests that V1 L2/3 excitatory neurons are equally able to contribute to detection regardless of their visual sensitivity, differential contribution based on projection target is still a possibility. This could be addressed in the future by retrogradely labeling various projection subclasses in V1, which may form anatomical and functional subnetworks ^65,66^. Examining projection-specific groups in deeper cortical layers, such as striatum or SC-projecting layer 5 neurons, would also be informative. Furthermore, our study argues that an unweighted readout is employed in L2/3 of the visual cortex for stimulus detection. It is possible that other cortical regions or layers use a weighted readout, as was suggested by an electrical nano-stimulation study in the rat barrel cortex^67^.

An important question for future work is how the timing of spikes in V1 impact visual detection. Although readout via GCaMP6 precludes the precise assignment of spike times, our photo-stimulation approach has sub-millisecond precision^24^. While previous work studying the detection of purely artificial optogenetic stimuli suggested that synchrony was not relevant for performance, readout strategies may differ between detection of direct neural stimulation and real visual stimuli^68^. Future experiments could probe the causal role of spike synchrony or different frequencies of induced neural activity by optogenetically controlling cortical ensembles with high temporal resolution photo-stimulation. The downstream decoder of activity might still use an estimate of the mean V1 activity, but the relevant timescale of spike count integration, the role of coincident or synchronous spiking (which could increase with greater mean activity), and whether the strategy evolves over the course of a detection trial remain to be determined. Future studies could also leverage faster GCaMP sensors or high-speed voltage imaging to examine if the timing of stimulation and the recurrent impacts on the activity of neighboring V1 neurons is important for detection.

Notably, a simple mean activity decoding strategy performs worse than the mouse on the task based on our imaging data. Superficially, this would seem to imply that mice must use a decoding strategy that performs better than a total mean decoder. However, this difference could be due to several technical factors related to measuring brain activity. One possibility is that other features of neural dynamics contribute to stimulus detection that none of our decoding models take into account (e.g., spike timing dynamics). Another possible explanation (which is not mutually exclusive) is that our imaging data does not accurately capture the mean of V1 of activity. It is currently nearly impossible to measure the activity of all individual V1 neurons. Additionally, GCaMP6 often poorly resolves low spike counts that may be induced by low contrast stimuli *in vivo*. These constraints exist for nearly all currently available observational approaches. Furthermore, we only measured activity in L2/3, while all layers of V1 will encode the stimulus. However, our highly specific and targeted causal perturbations were able to adjudicate between the weighted and unweighted decoding strategies. This demonstrates the advantage of such causal perturbations over relying purely on a correlation between the performance of various decoding models and the animals’ performance on the task.

A previous study of photostimulation in visual detection, found that behavioral performance and the impact of photostimulation depended on internal state, as measured by pupil size and pretrial population correlations^34^. Another study of detection of artificial stimulation in the barrel cortex found that pretrial correlations, but not pupil size, correlated with artificial detection^31^. Other studies in similar tasks have found mixed results of whether pupil size or other internal state metrics correlate with behavioral performance^69–72^. In our task, we found no relationship between pretrial pupil size or population correlations on behavioral performance (Fig. S9A-B,D). We also saw no evidence that photostimulation might alter behavioral state (Fig. S9C). Despite this difference, the prior study in visual detection also showed that groups of ∼20 neurons could impact behavior under engaged conditions, in agreement with our results^34^. However, the authors only stimulated co-tuned, visually responsive neurons and therefore did not address whether the co-tuning or the visual sensitivity of the targeted ensemble was in fact required to alter behavior. Further investigation to determine how behavioral state and network influence interact would further illuminate the mechanisms by which ensembles contribute to perception.

While we suggest that a weighted readout of neurons with different stimulus tuning within a retinotopically aligned region is not the strategy for detection, the mouse may still perform a “weighted” readout of different areas of V1 across the retinotopic map. In fact, previous studies have shown that mice can direct spatial attention to specific regions and that retinotopic alignment of silencing is key for its effects^18,61,73^. It would be interesting for a future study to photostimulate retinotopically misaligned regions and determine how these can impact detection if at all.

In our study, photostimulation of any ensemble, regardless of its properties, did not result in an increase in false alarms. This contrasts with prior work showing that electrical microstimulation or one photon optogenetic stimulation can induce behavioral responses in detection tasks in the total absence of a stimulus (and can induce phosphenes in humans)^23–25^. We speculate that the relatively small number of spikes we added to the network in this study (most likely adding on the order of 3-4 spikes per neuron across 10-30 neurons)^50^, is simply insufficient to drive artificial perceptions. Our goal was to determine how different patterns of neural activity could boost detection of real stimuli, and our relatively focal and mild augmentation of neural activity was sufficient to address this question. Larger scale targeted holographic stimulation can co-activate hundreds^33,50^ of unique neurons, and leveraging such a photo-stimulation regime should be useful for identifying the optimal stimulation scheme for generating false perception, as might be useful for neural prostheses.

In this study we focused on stimulus detection – a necessary step in sensory perception that precedes stimulus identification. Notably, for some types of sensory discrimination (e.g., between two stimuli with identical contrast but different orientations) a mean activity decoding strategy could not support task performance. In such tasks, a comparison between population activity patterns, and possibly the weighting of specific cortical ensembles, is likely to be required. We speculate that the brain uses a simple mean decoding strategy for detection, despite its sub-optimality, because it may promote generalization. Any stimulus that increases mean V1 activity can be decoded with this strategy, simplifying the readout. Once a stimulus is positively detected, we surmise that the cortex may then switch to a different readout strategy to promote stimulus identification. Indeed, in the visual and somatosensory systems a transition occurs between early, non-adapted stimulus representations that are more detectable but less discriminable, and late, adapted representations that are less detectable but more discriminable, suggesting that such a mode switch could occur^74,75^. Analogous experiments to those we performed here, but in a discrimination task, can be used to address the causal features of neural codes that enable stimulus identification. Related work in mouse V1 implies that photo-activating specific cortical ensembles that preferentially encode certain orientations can contribute to performance in an orientation discrimination task^32,33^. However, similar studies in mouse barrel cortex in animals performing tactile discrimination tasks have yielded conflicting findings^46,76^. These latter studies have instead highlighted either the choice encoding of the targeted ensemble, or simply the total number of photostimulated neurons – even of those that do not appear to encode the discriminated stimuli (similar to our results). In the latter case, the animals had to discriminate which side of the face received stronger whisker stimulation. In this task mice might also have employed a simple cortical activity summation strategy because a comparison of total neural activity between the two hemispheres could be a viable strategy to solve the task, although neurons with tuning to both sides existed in both hemispheres. This suggests that unweighted activity read out may be employed in many different types of tasks. Additional work is needed to determine if and when the cortex uses weighted decoding strategies in discrimination and resolve these apparent conflicts.

In sum, our results suggest a simple model in which visual stimulus detection relies on a general readout of V1 neural activity without preferentially weighting the activity of more visual sensitive neurons. This strategy is suboptimal in the specific conditions of our stereotyped detection task, but may promote generalization to many different stimuli at the expense of optimal performance for a particular stimulus class. These surprising results suggest that the suboptimal performance that is frequently observed across species, modalities, and tasks^3,4,7,11,19,49,54,59^ may not result from readout noise in downstream regions but from suboptimal readout strategies. These may include a generalized unweighted readout strategy in detection, as we showed here, to shorter-than optimal integration windows in motion discrimination^77,78^ in monkeys, to ignoring anti-tuned neurons to inform orientation discrimination^59^ in mice, to location-invariant coding of object recognition even when stimulus location is stable in a task, as has been suggested to account for suboptimal human performance^79^. This may reflect that optimal performance across the wide variety of demands that natural conditions present may be computationally infeasible^80^. Considering a behavior task within its ethological contexts may reveal that a putatively “suboptimal” strategy is in fact preferable in more variable natural conditions.

## Acknowledgements

We thank M. Wang and D. Quintana for assistance with behavioral training and K. Gopakumar, S. Sridharan, and J. Beyer for scientific and logistical support. Additionally, we thank the members of the Adesnik lab for comments on the manuscript.. This work was funded by NEI grant R01EY023756 and RF1NS128772, the National Science Foundation Graduate Research Fellowship DGE-1752814, and a fellowship from the Chan-Zuckerberg Biohub.

## Author Contributions

H.A. and H.A.B. conceived of the study. H.A.B. designed and performed all experiments and analysis with advice from H.A. H.A.B. and H.A. wrote and edited the paper.

## Methods

### Mice

All experiments were approved by UC Berkeley Animal Care and Use Committee. Mice (c57/bl6) used were double heterozygous CaMK2-tTa; tetO-GCaMP6s mice of both sexes (6 female, 8 male). Some mice additionally carried one of Vglut1-IRES-Cre, PV-Cre, Emx-Cre, or SST-Cre, which were not relevant to the experiment as all gene expression (both viral ChroME2S and transgenic GCaMP) was driven by the TRE promoter and thus was tTa-dependent, not Cre dependent. Mice were aged 12-47 weeks.

### Microscope Design

2-photon optogenetics was accomplished using (3D-SHOT)^81,82^. Briefly, the microscope was custom-built on the Movable Objective Microscope (MOM; Sutter Instrument Co.) platform. A 3D two-photon (2p) photostimulation path with a Monaco 1035-80-40 (1035 nm, 2 MHz, 276 fs, Coherent Inc.) and a fast resonant-galvo raster scanning 2p imaging path with a Chameleon Ultra Ti:sapphire laser (Coherent Inc.) were merged by a polarizing beamsplitter before the microscope tube lens and objective. In the 3D-SHOT photostimulation path, a custom temporally focused pattern (CTFP) was by a blazed diffraction grating (33010FL01-520R Newport Corporation), which was then relayed through a rotating diffuser to randomize the phase pattern and expand the beam to cover the area of the spatial light modulator (SLM; HSP1920-1064-HSP8-HB, 1920 × 1152 pixels, Meadowlark Optics), which replicated the CTFP in 3D using holographic phase masks. Holographic phase masks were calculated using an iterative implantation of the Gerchberg-Saxton algorithm^90^. An Arduino Mega (Arduino) was use to gate the photostimulation laser to only be on during the turn phase of the resonance galvos bidirectional line scan, in order to restrict photostimulation artifacts contaminating the imaging data to only the edge of the FOV, which was not analyzed. Multiplane imaging was accomplished using an electrically-tunable lens (Optotune) placed in the light path prior to the resonance scanners. The two paths were calibrated as described previously^43^.

### Viral injection, headpost implantation, and cranial window

Mice 8 weeks or older were anesthetized with 2% isoflurane and administered 2 mg/kg of dexamethasone to reduce swelling and 0.5 mg/kg of buprenorphine for analgesia. Mice were secured in a stereotaxic frame with a heating pad. An incision was made to expose the skull, which was then covered with Vetbond (3M). Mice were then intracranially injected with PhP.eB-TRE-ChroME-P2A-H2B-mRuby3. 4 burr holes were placed over left V1 (2.7 mm lateral, 1 mm anterior of lambda) approximately 0.75 mm apart. 125 nL of virus was injected at 50 nL/minute at two depths: ∼350 µm and ∼250 µm below the dura. This was done with a microinjector (Micro4; World Precision Instruments) and a glass pipette (Drummond Scientific). The glass pipette was kept in place for 5 minutes after injection.

A craniotomy was made over the injection sites and V1 using a 3.5 mm biopsy punch. Once the skull was removed, cold PBS was used to flush the area, and then a cranial window was placed. Cranial windows consisted of two 3 mm diameter circular coverslips and one 5 mm diameter circular coverslip glued with optical glue. Metabond (C&B) was used to secure the cranial window. Finally, a custom-made titanium headplate was attached using Metabond (C&B) and OrthoJet (Lang Dental).

### Imaging and Photostimulation during Behavior

Mice were head-fixed under the objective in a custom-designed tube with paw rests. Stimuli were presented on a 2048 x 1536 Retina iPad LCD display (Adafruit Industries), with the backlight LED controlled to turn on only during the turn phase of the imaging galvos to avoid artifacts in the imaging. In a subset of recordings, pupil diameter was monitored on the ipsilateral eye (not facing the monitor). The shutter of the camera was synced to the frame acquisitions in ScanImage, such that the framerate was the volumetric imaging rate times the number of planes. The camera imaged IR light exiting the pupil from the imaging laser.

Fast 3 z-plane volumetric imaging at 5.88-6.36 Hz was conducted (field-of-view: 821 x 761) using ∼75mW of imaging power at 920 nm. A 20-25 ms delay was added to the y galvo time to allow the Optotune (see microscope design) to settle. Planes were 30 µm apart.

At the start of the session, the retinotopic preference of the field of view was mapped using custom MATLAB code and Psychophysics ToolBox. Small (12°) drifting gratings (2 Hz, spatial frequency: 0.08 cpd) that iterated through 8 directions. Stimuli lasted 2 seconds with a 2 second inter trial interval. Stimulus placement for the rest of the experiment was determined by online monitoring of the preferred retinotopic location of the field of view, based on average fluorescence responses of the whole field.

In 2-photon stimulation experiments, two additional epochs preceded the final behavioral testing. First, a short (∼75-100 trial) behavioral session with only high contrast stimuli (64% and 100% contrast, otherwise identical to description below) was presented to determine the task responses of neurons. Then, the retinotopic mapping and behavioral session were analyzed in suite2p^83^.

During suite2p analysis, an image of the field of view was taken at 1040 nm to visualize the red nuclei of opsin-expressing cells. Cell locations were determined using either a custom peak-find algorithm in MATLAB or Cellpose nuclear detection^84^. Then all cells in the field of view were stimulated sequentially in groups of 5-15 with 5 pulses of 5 ms duration, typically at 2 different powers.

Finally, the online visual response properties (analyzed using suite2p) and photostimulation response properties (analyzed using scanimage online roi analysis) were matched by cell location and used to determine 1-2 groups of neurons to stimulate. Online photostimmability was determined as a significant (p<0.05, t-test) response to photostimulation versus no photostimulation at the desired power. In some sessions, neurons were chosen by photostimulation response alone (random). In others, two groups of photostimmable cells were chosen, one that was significantly responsive to visual stimuli (p<0.05, either one-tailed paired t-test for pre versus post response to visual stimulus trials or one-tailed independent t-test for 100% contrast visual stimulus versus no contrast trials) for in the short behavior epoch, and one that was not. These criteria were only for online stimulation selection, and were not used for any analysis presented in this paper. The pre-test photostimulation and short behavioral epoch data is also not included here. For the purposes of analysis, all groups were classified post-hoc regardless of the online determination.

### Behavioral testing

Behavior was controlled by a custom-written program in Psychopy^85^ using National Instruments Data Acquisition (NI-DAQmx) to monitor licks and dispense water. Visual stimuli consisted of 20 degree static visual gratings of the same orientation and phase, with spatial frequency .08 cycles per degree. For the behavior, stimuli were displayed for 500 ms, followed by a 1000 ms response window. During go trials (contrast > 0%), mice received a water reward for licking in the response window, with no reward or punishment for licking during the display period. Intertrial intervals were 3-9 seconds. Mice received time outs (4-9 seconds) for licking during the response window of catch trials or during intertrial intervals (following a 1 second grace period). Licks were detected with a custom-designed optical lickport using an infrared beam break system. A 200 ms delay occurred between trial initiation and visual stimulus presentation to ensure all adequate time for the SLM to load photostimulation conditions, and during this time timeouts could not be delivered. Any trials with licks in this 200 ms pre-trial time or within the first 150 ms of visual stimulus presentation were excluded from analysis.

### Photostimulation in behavior

In sessions with photostimulation, trials were equally divided between each photostimulation condition (including no stimulation). Either one or two groups were photostimulated. Mice were rewarded or punished according to the visual stimulus condition of the trial, regardless of photostimulation.

Neurons in each photostimulation group were stimulated simultaneously in one hologram. Only one group per trial was photostimulated. Power was selected per field of view and was uniform (after correction for diffraction efficiency of the SLM) across all neurons. Neurons were stimulated with three 5-ms long pulses at either 20 or 30 Hz, beginning 35 ms after visual stimulus onset, approximately the time the first visually-evoked spikes reach V1^17,39^.

In some sessions, additional photostimulation or control trials with no visual stimuli were presented during long intertrial intervals to collect more data about the recurrent effects of 2-photon stimulation groups. These trials did not change any timing rules of the task. These trials were not included in any behavioral analysis, but were included in neural analysis and decoding analysis. Any of these interbehavioral trials with licking at any time within the 1.5 second long trial were excluded.

In all sessions, imaging was continuously monitored and spatial drift corrected manually.

### Behavioral Training

Behavioral training was as described previously in raspberry pi controlled boxes using drifting gratings of the same orientation and spatial frequency as final testing^24,25,44^. After learning the task outside of the microscope, mice were habituated to the microscope rig, then trained on the microscope with static gratings until reaching satisfactory performance for testing.

### PPSF (physiological point spread function) Experiments

Cells were first screened for photostimmability as described in above. Then two groups of neurons (6-11 neurons each) were selected for physiological point spread experiments. Cells were stimulated at their detected center, and at 12 points radially offset between (−3 and 30 μm) and 10 points axially offset between −6 and 48 μm. Axial offsets that exceeded the calibrated area of the microscope were eliminated, which occurred for neurons on upper planes at the large axial offsets. For analysis, axial and radial point spread functions were fit with a gaussian function. To be included, neurons had to be significantly photostimmable and both the radial and axial fit had to have r^2^>0.1.

### Analysis

#### Calcium imaging analysis

All calcium data was extracted and neuropil subtracted using suite2p ^83^ with manual curation of cell identity. Z-scored ΔF/F was calculated by first using a rolling 20th percentile estimation of F_0_ to calculate ΔF/F, then z-scoring the traces over the entire recording. Trials were synchronized to a stimulus onset signal sent by the Psychopy behavioral system.

Activity was analyzed in the 666 ms after stimulus onset. Unless specified otherwise, photostimulation population activity and activity changes were measured in no-visual-stimulus and no-lick trials to isolate recurrent impacts and eliminate any confounds of photostimulation-induced changes in behavior. Visual and choice response properties were measured in no-photostimulation trials.

#### Inclusion criteria

To be included for any analysis, sessions had to have at least 15% of neurons significantly positively visually responsive. Fewer than 25% of trials had to be excluded for motion (see below). The false alarm rate for no photostimulation conditions had to be below 33%. At least 90 trials per photostimulation condition had to be included (see criteria below). Finally, at least 75 trials for each photostimulation condition had to fall within periods where the mouse had high performance (defined as responding at least 60% of the time to no-photostimulation 100% contrast trials for at least 50 trials in a row). This criteria was used to ensure a sufficient portion of the session had good behavioral performance, but trials outside of these windows were included in the analysis.

To be included for analysis of photostimulation effects, sessions had to have at least 50% of targeted neurons (defined as the closest coplanar neuron within 13 μm of the holographic spot) significantly activated by photostimulation

Trials were excluded if they had trials with licks in the 200 ms pre-trial time or within the first 150 ms of visual stimulus presentation. Trials with where any frames in the 1 second pre-stimulus or two seconds post-stimulus period were marked as “bad frames” by suite2p, or where the mean euclidean displacement of the field of view (as determined by suite2p motion correction) from 166 ms pre-stimulus onset to 666 ms post-onset was greater than 6.5 μm were also excluded, to avoid errors in holographic targeting from brain motion. For behavioral analysis but not neural analysis, behavioral sessions were cut to include only periods where the mouse maintained basic attention criteria (was responding to trials with a visual stimulus > 8% contrast at least 15% of the time (in a rolling window of 10 trials)) for at least 50 trials. This typically removed only a small fraction of trials, if any.

#### Visual sensitivity and choice selectivity

Visual sensitivity was defined as the area under the roc curve for AUROC for visual stimulus presence for all trials (regardless of lick response). Bootstrapped 95% confidence intervals were constructed by resampling the data and used to determine significance of sensitivity. For analysis of significantly visually responsive neurons in Fig. 3B, S5 & S5 significance was determined by comparison of mean activity 5 frames pre-visual stimulus to 5 frames post-visual stimulus (p<0.01, one-tailed Wilcoxon signed-rank). Choice selectivity, also called choice probability, was measured in only no-photostimulation, visual stimulus trials, and only at contrasts where the animal was performing at least 10% better than the false alarm rate at 10% worse than peak performance, to ensure an adequate number of hit and miss trials to measure choice. We used the approach described in Kang et al 2012^55^ in order perform balanced z-scores and combine data across contrasts. Finally, this z-scored activity was used to calculate the AUROC for the animal’s choice.

#### Photostimulation populations

Neurons were considered directly activated if they were significantly activated by 2-photon stimulation in no visual stimulus trials, comparing mean activity 5 frames pre-visual stimulus to 5 frames post-visual stimulus (p<0.01, one-tailed Wilcoxon signed-rank test) and within 21 μm radially (in xy) of a target spot, regardless of their axial distance, a conservative determination made based on physiological point spread function measurements (Fig. 2B, S1). We always considered the activity of all directly activated neurons, including random off-targets, to understand the full effect of photostimulation. The indirect population was defined as neurons further than 21 μm radially from a holographic spot.

#### Ensemble classification

Ensembles were considered network-activating if the mean population activity on photostimulation trials was greater than ⅓ of a standard deviation above the response in no photostimulation trials. To determine if an ensemble was visually sensitive, we considered the mean visual sensitivity of the ensemble. Then, we drew 1000 random sets of neurons, matched in size to the directly activated ensemble and determined the mean and standard deviation of the mean visual selectivity of these randomly drawn ensembles. Ensembles were considered sensitive if they had a visual sensitivity greater than one standard deviation above the mean of the random draw.

#### Decoding analysis

Both the mean activity decoder and weighted decoder used the python sklearn package LogisticRegression class using the “lbfgs” solver, “l2” normalization, and an inverse regularization strength of 0.001 (the latter two values do not matter for the mean activity decoder as there is only one input). Both were trained to report stimulus presence across all presented contrasts in no photostimulation trials. Models were trained and assessed on fifteen repeats of a five-fold stratified cross-validation that fairly represented all presented contrasts. The mean decoder was fed the mean activity of all imaged neurons in the analysis window for each decoder. The weighted decoder was fed the mean trialwise activity of all imaged neurons separately.

Decoder AUROC at each contrast was calculated by the probability output of the decoder for the held-out trials at that contrast versus held-out no stimulus trials. For comparison with mouse behavior, discriminability index was calculated as 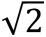 ∗ 𝑍(𝐴𝑈𝑅𝑂𝐶)^86^, where Z is the inverse CDF of the normal distribution.

For comparison, we performed the same analysis as above, but used a support vector classifier with a sigmoid kernel (sklearn SVC, inverse regularization strength 2.5 for weighted and 1 for unweighted) and a linear support vector classifier (sklearn LinearSVC with an “l2” normalization and inverse regularization strength of 0.001).

#### Behavioral analysis

% change in hit rate was calculated as the difference in mean hit rate at test contrasts for photostimulus vs no photostimulus trials, divided by the mean hit rate at test contrasts for no photostimulus trials. Boostrapped 95% confidence intervals were calculated by resampling photostimulus trials 50000 times with replacement. For display purposes, psychometric curves were fit with a modified weibull function.

#### Pretrial correlation analysis

For the analysis in Fig. S9D, we took the 3 seconds of pretrial activity and computed the pairwise Pearson correlation coefficient between the activity of all possible neuron pairs, and then took the mean of all coefficients, as done previously^34^. We did this for both deconvolved traces (using suite2p deconvolution) and the z-scored DFF.

#### Pupil analysis

Pupil area was quantified using a modified version of the ellipse-based algorithm of FaceMap^87^, tuned to fit the needs of imaging a light pupil on a dark background and remove reflections of the laser light. n=19/31 sessions had pupil image performed, 3 had to be excluded due to oversaturated camera gain or bad camera angle causing the eyelid to occlude too much of the pupil. Within the usable sessions, we excluded any frames where the pupil eccentricity was more than 2x the standard deviation over the mean of the pupil eccentricity. Then, we calculated normed pupil area by subtracting the 5th percentile of pupil area and then dividing by the 90th percentile pupil area. To quantify pretrial pupil relationship to behavior, we took the mean normed pupil area in the 1 second pre stimulus onset for all miss and hit trials at test contrasts (contrasts between 1-99%). Only no-photostimulation data was included in this dataset. Then, to examine the impact of photostimulation on pupil size, we looked at only no-visual stimulus presentation, no-response trials an considered activity in the 1 second after photostimulation offset.

---

**Figure S1:**
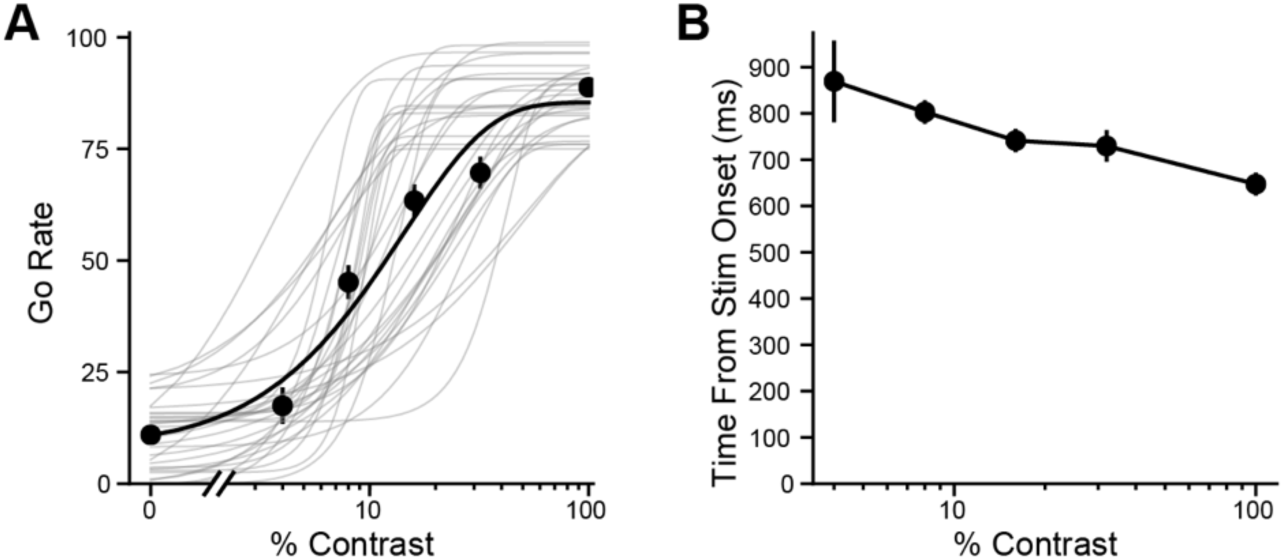
Summary of behavioral data. **A:** All fit psychometric curves (gray) and mean (black) for n=31 sessions used in behavioral analysis. **B:** Average lick response time from stimulus onset for go trials. Stimulus ends at 500 ms. Data are plotted as mean ± sem.

**Figure S2:**
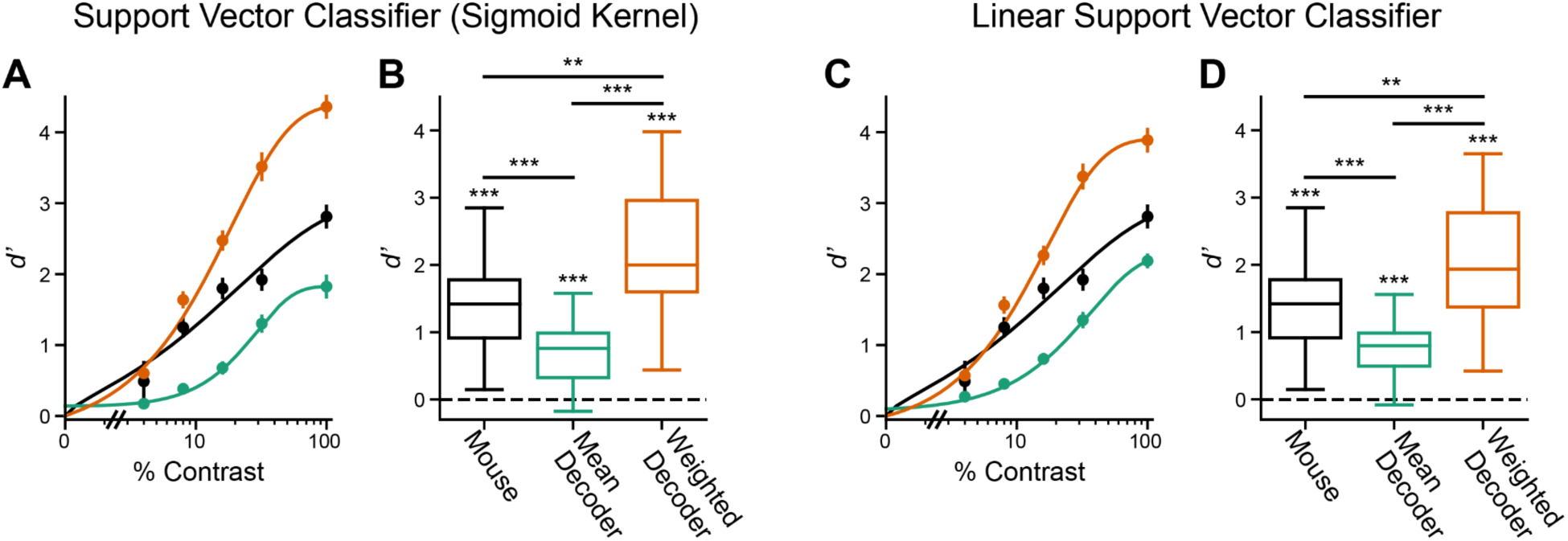
Multiple classifier types show stimulus decoding for both weighted and mean decoders. **A-B:** Same as Figure 1D-E but for a support vector classifier with a sigmoid kernel. **A:** Comparison of the cross-validated *d’* (discriminability index) of mouse behavior, and decoders trained to detect stimulus presence. Teal, a decoder on mean activity. Orange, a decoder allowed to vary the weights of individual neurons. **B:** Comparison of cross-validated *d’* of different models tested at test contrasts (contrasts between 1-99%). mouse: p=7.0e-6, mean decoder p=2.2e-5, weighted decoder: p=7.0e-6. Between models: mouse vs mean decoder p=2.3e-4, mouse vs weighted decoder p=9.0e-3, mean decoder vs weighted decoder p=7.0e-6, multiple-comparisons corrected (mc) Wilcoxon signed rank. **C-D:** As in A-B but for a linear support vector classifier. **C:** Comparisons vs chance: mouse: p=7.0e-6, mean decoder: p=8.6e-6, weighted decoder: p=7.0e-6. Comparisons between: mouse vs mean decoder p=3.5e-4, mouse vs weighted decoder p=2.2e-2, mean decoder vs weighted decoder p=7.0e-6, mc-Wilcoxon signed rank. **A&C:** Data plotted as mean ± sem in A&B.

**Figure S3:**
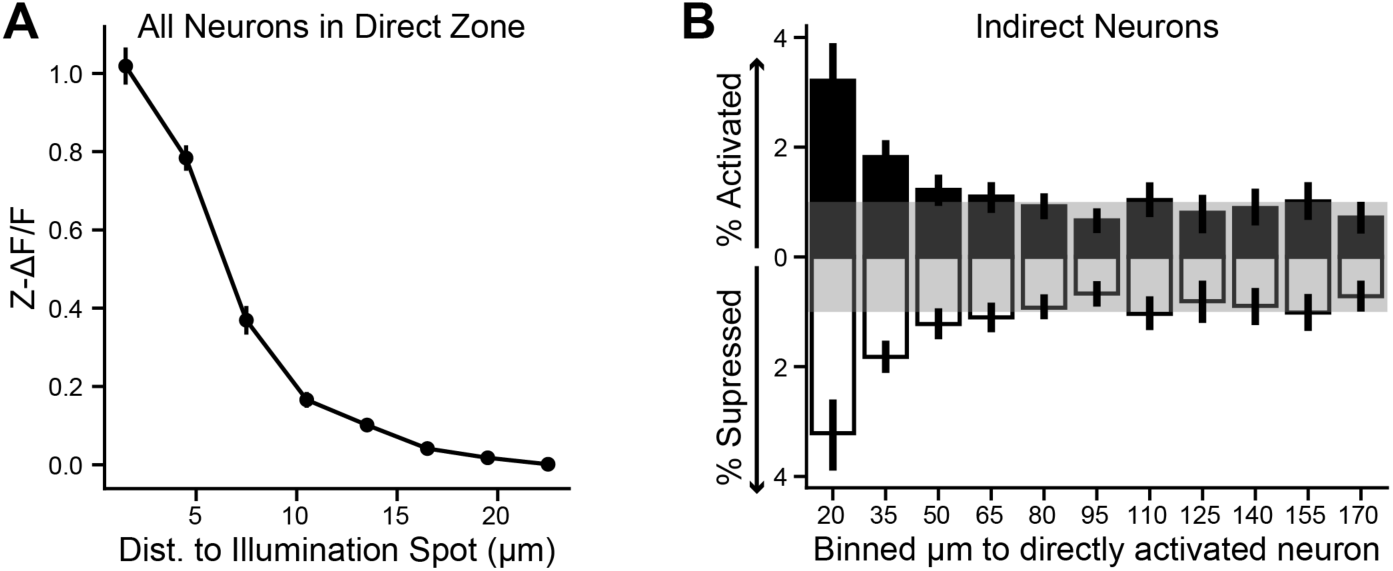
Analysis of spatial footprint of 2-photon stimulation. **A:** Activity evoked by 2-photon stimulation for all neurons within the direct zone (20 um from a holographic spot) versus distance to spot. **B:** For indirect neurons, % of neurons for each ensemble (n=41) that were significantly activated (black bars) or suppressed (white bars) based on binned distance from the nearest directly activated neuron. Gray shaded region indicates percentage expected by chance.

**Figure S4:**
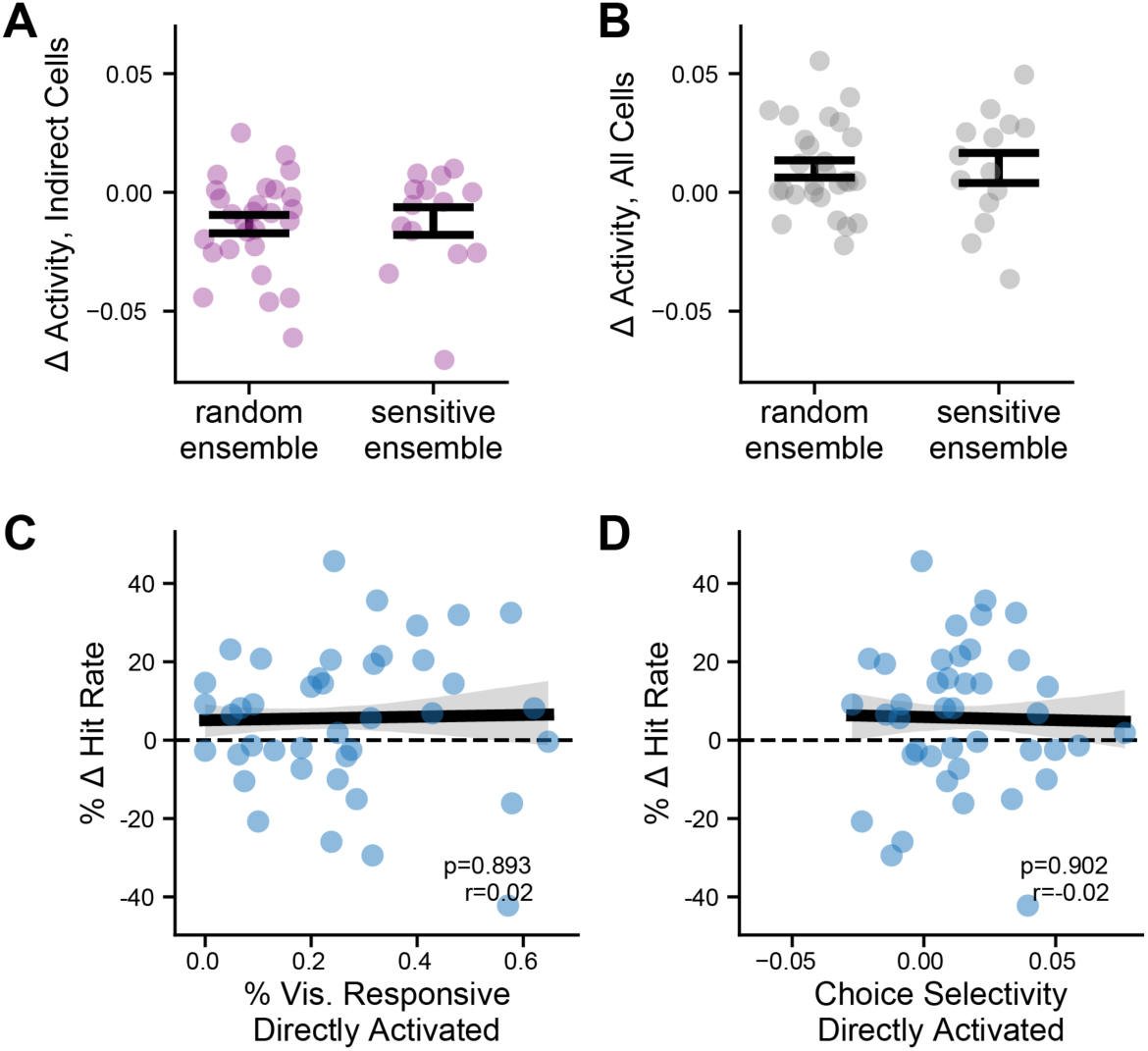
No relationship between visual and task response properties of ensemble and network or behavioral effects. **A-B:** No difference in indirect network activation (A) or total network activity change (B) between random and sensitive ensembles, indirect: p=0.62, all: p=0.74, Wilcoxon rank-sums test. **C:** % of directly activated neurons that are significantly visually responsive versus change in hit rate. **D:** same as A but for the mean choice selectivity (AUC) of directly activated neurons. Gray areas, 68% confidence intervals of regression. Statistics are for Pearson correlation coefficient.

**Figure S5:**
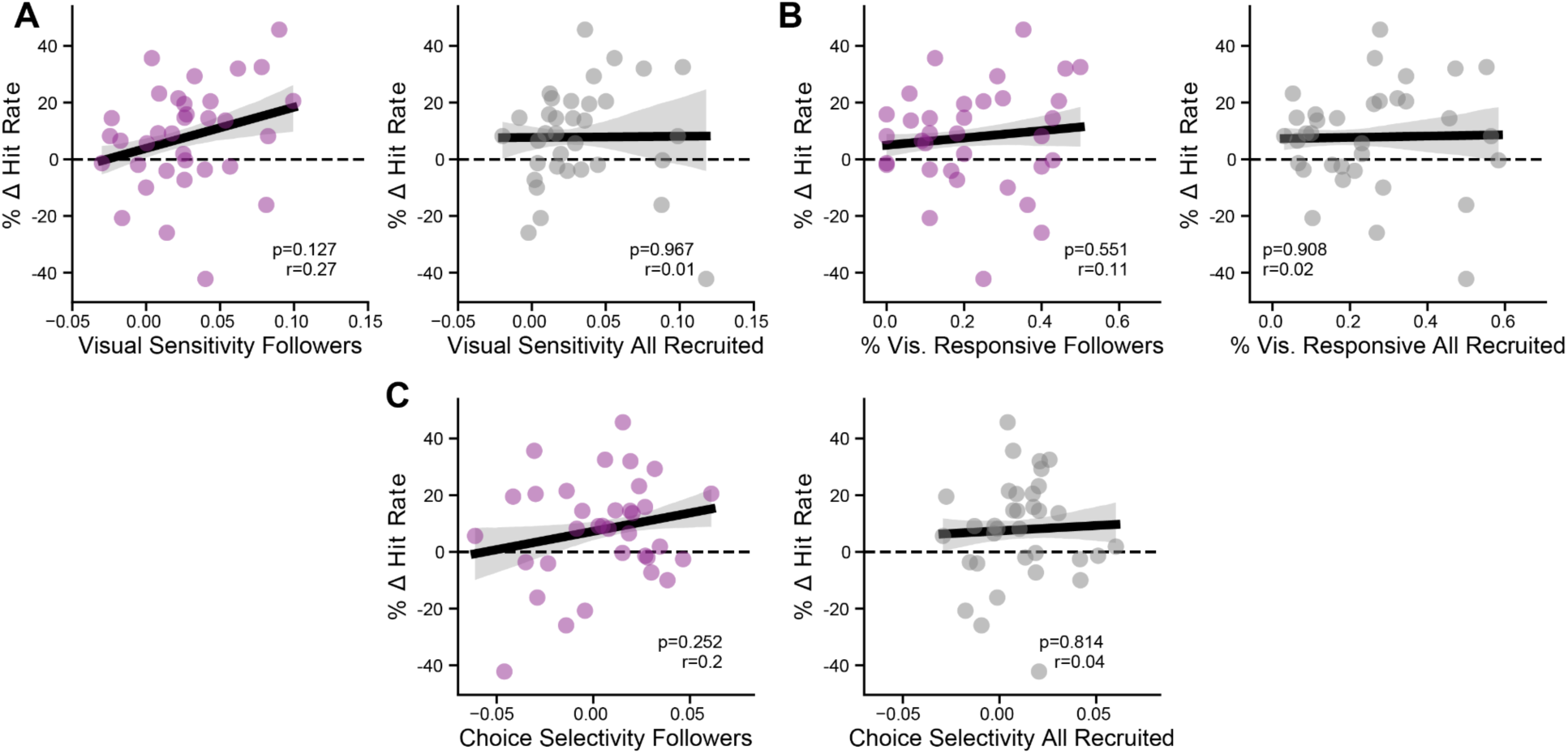
The visual properties of synaptically recruited neurons does not predict their behavioral impact. **A:** Mean visual sensitivity (AUC) for followers (left) and all recruited (right) versus % change in hit rate. Follower cells are indirect cells that are significantly activated by 2-photon stimulation, presumably by synaptic effects. All recruited neurons are the combination of directly activated and follower cells. For follower analysis only ensembles with at least four follower neurons (n=34) were used, all ensembles (n=41) were used. **B:** Same as A, but for % of significantly visually responsive neurons. **C:** Same as B, but for choice selectivity (AUC). **A-C:** Gray areas, 68% confidence intervals of regression. Statistics are for pearson correlation coefficient.

**Figure S6:**
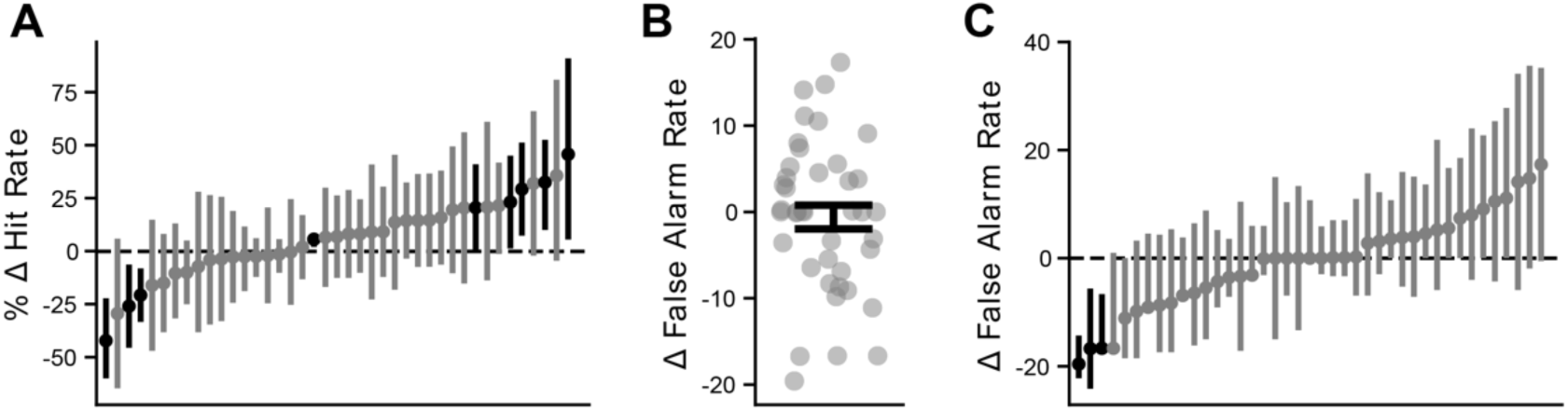
2-photon stimulation impacts hit rate but not false alarm rate. **A:** for each stimulated ensemble, bootstrapped 95% confidence intervals on the % change in the hit rate at test contrasts. Black lines indicated significantly different ensembles (9/41). **B:** Change in hit rate induced by 2-photon stimulation. p=0.78, Wilcoxon signed-rank test. **C:** Same as A, but for change in false alarm rate.

**Figure S7:**
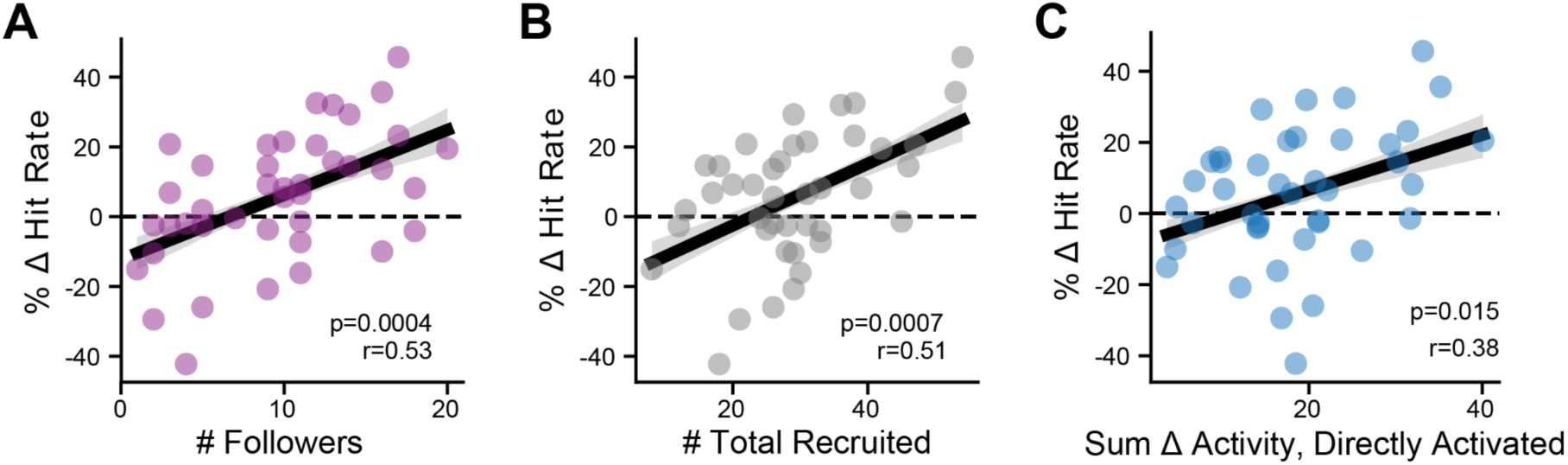
The number of recruited neurons and strength of direct photostimulation predicts behavioral outcome. **A:** Number of recruited follower cells (indirect cells that are significantly activated by 2-photon stimulation) versus % change in hit rate(n=34 ensembles with at least four followers). **B:** Same as A but for total number of recruited neurons (directly activated and follower) (n=34 ensembles with at least four followers). **C:** The sum of change of activity across all directly activated neurons. Taking the sum, rather than mean, activity change is important because of the small number of directly photostimulated neurons and the variability in their number across sessions. n=41. Gray areas, 68% confidence intervals of regression. Statistics are for pearson correlation coefficient.

**Figure S8:**
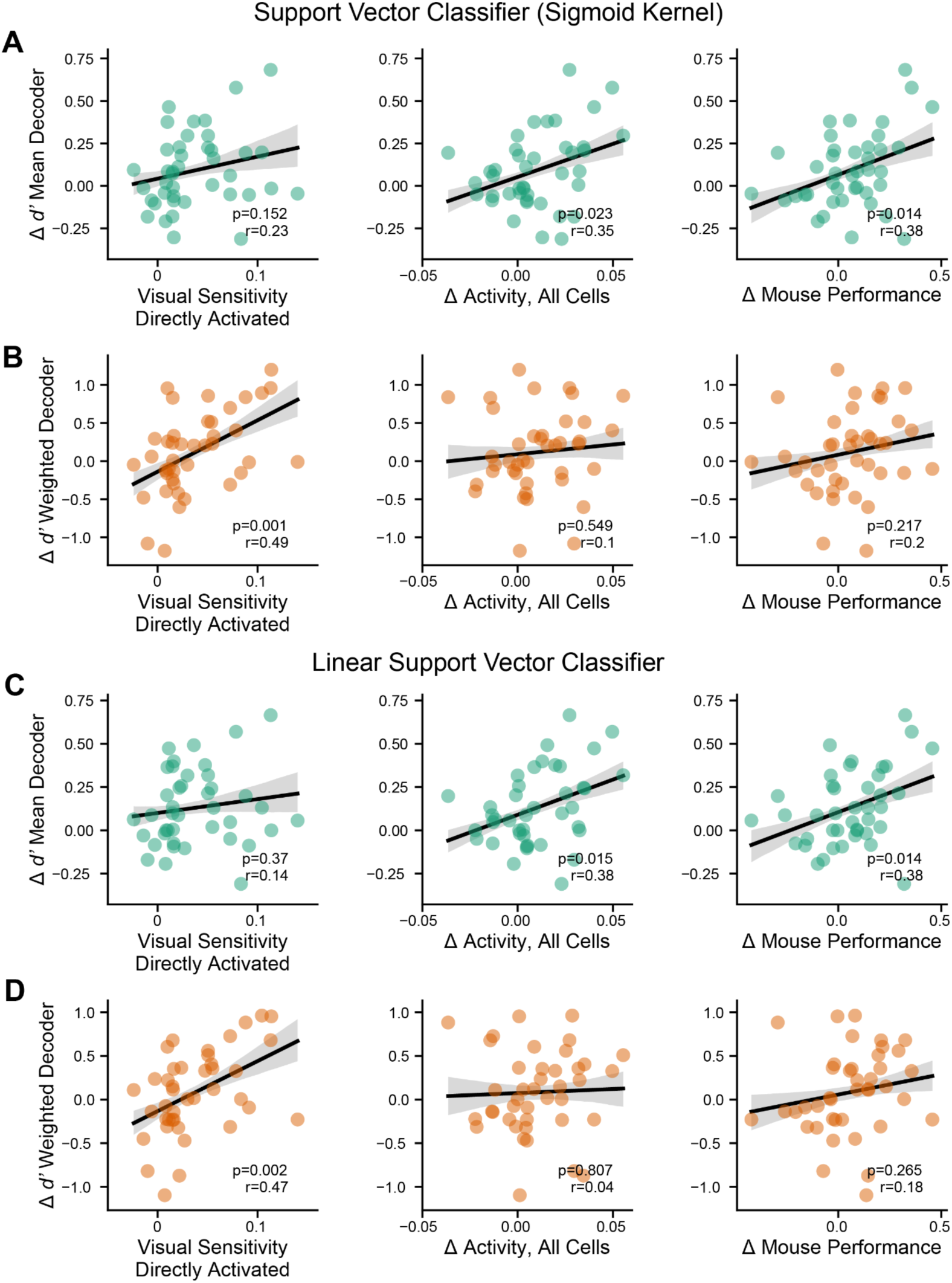
2-photon stimulation induced changes in a mean decoder are more consistent with behavioral changes. **A:** For a support vector classifier with a sigmoid kernel (SVC), the change in mean activity decoder discriminability index between 2-photon stimulation and no stimulation trials at test contrasts compared to average visual sensitivity (AUC) of directly activated neurons (left), photostimulation-induced change in activity of all neurons (center), or mouse’s change in hit rate (right). Gray area, 68% confidence interval of regression. Statistics are Pearson’s correlation coefficient. **B:** Same as A, but for weighted SVC. **C-D:** Same as A-B but for a linear support vector classifier (LSVC).

**Figure S9:**
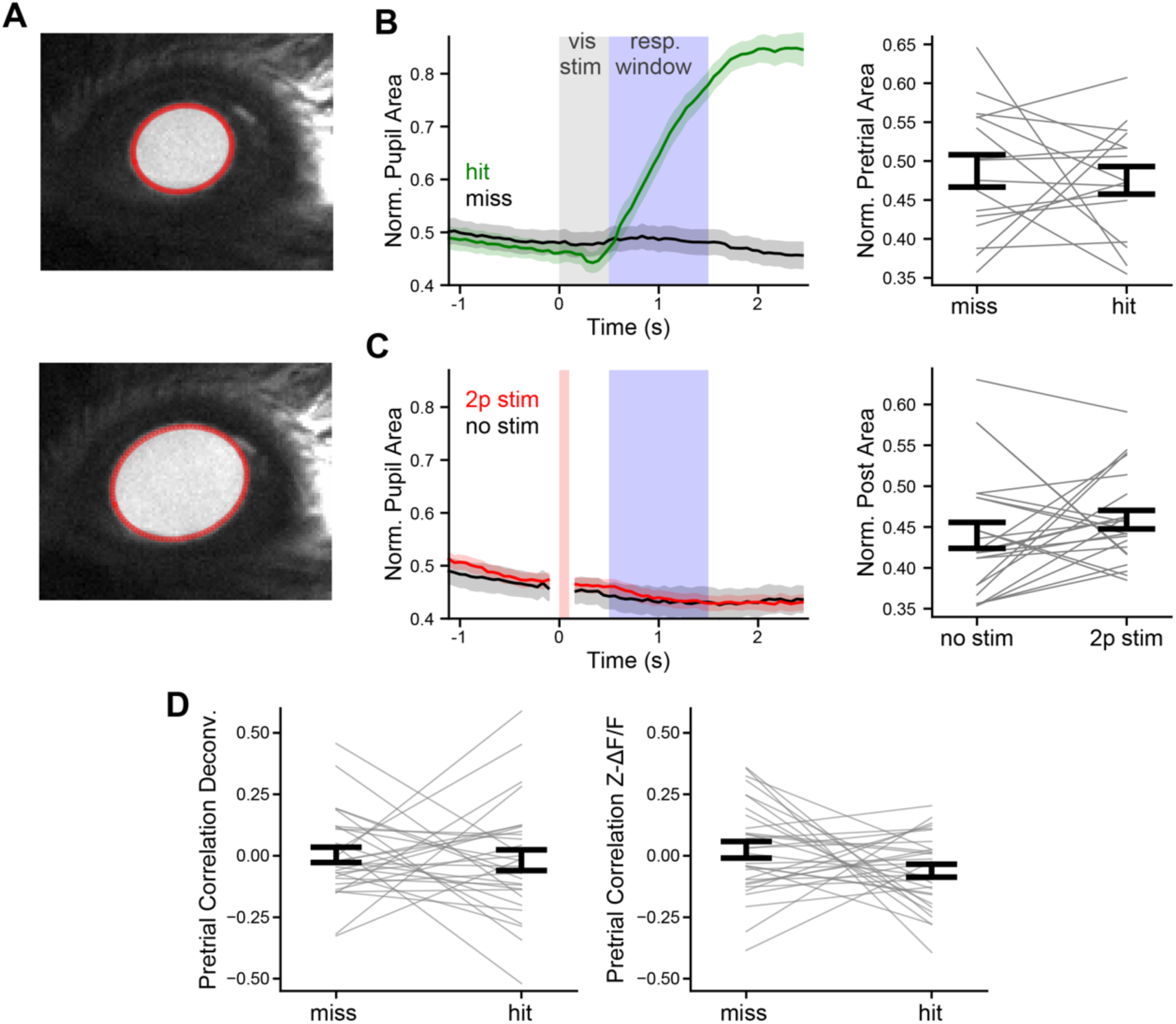
Pretrial pupil area and neural synchrony do not predict behavioral outcome. **A:** Example images of pupil tracking for a small pupil area (top) and large pupil area (bottom). Red, tracked pupil area. **B:** Left, average timecourse of pupil size for all miss and hit trials across n=16 sessions. Right, comparison of pupil area averaged over 1 second before stimulus presentation for hit versus miss trials. p=0.92. n=16 behavioral sessions. **C:** Analyzing the effect of 2-photon stimulation on pupil size in no visual stimulus, no response trials. Left, average time course of pupil size between 2p stim and no stim trials. Data during 2-photon stimulation is excluded due to contamination of pupil tracking by the stimulation laser light. Right, average pupil area for the one second post stimulation time between stimulation and no stimulation trials p=0.29, Wilcoxon signed-rank, n=23 photostimulated ensembles. **D:** Pretrial correlation, measured as the z-scored trialwise mean pairwise correlation of all neurons, compared between miss and hit trials at test contrasts. Left, measured from deconvolved calcium traces. p=0.43, Wilcoxon signed-rank, n=31 behavioral sessions. Right, measured from z-scored DFF p=0.22, Wilcoxon signed-rank, n=31 behavioral sessions.

